# Microbial Scl1 Activates TGFβR1 receptor kinase signalling to Drive Fibrosis – Inflammation Axis in Alcohol-Associated Liver Disease

**DOI:** 10.1101/2025.07.08.663677

**Authors:** Manisha Yadav, Gaurav Tripathi, Anita Nandi, Shabnam Shabnam, Jayesh Kumar Sewak, Vasundhra Bindal, Neha Sharma, Sanju Yadav, Vipul Sharma, Nupur Sharma, Babu Mathew, Yash Magar, Deepanshu Deepanshu, Sushmita Pandey, Rimsha Saif, Sanjeev Das, Abhishak Gupta, Anupama Kumari, Devender Seghal, Shvetank Sharma, Shiv Kumar Sarin, Jaswinder Singh Maras

## Abstract

**Background and Aims:** Alcohol consumption alters gut microbiota, which can affect metabolism, immune regulation, and signalling pathways and lead to alcohol-associated liver disease (ALD). We investigated how alcohol-associated gut microbiota (AGMs) modulate liver kinome signalling to drive inflammation and fibrosis in ALD.

**Method:** Liver kinome and metaproteome changes were studied in rats (n=6/group) colonized with stool of severe alcohol-related hepatitis (SAH) patients (SAH→healthy-rats) or healthy human donors (HD→ALD-rats). AGM-associated liver kinome changes were cross-correlated with bacterial genera. A bacterial protein; Streptococcal collagen-like protein 1 (Scl1) was identified with affinity for TGFβR1. This was validated by molecular docking and immunoprecipitation-LCMS assay. Expression of *Streptococcus pneumonia* and *pyogenes* was done in stool of SAH patients (n=10). Also, Scl1 was quantified in patient stool (n=24) and liver tissue (n=19) samples. Validation was performed by assessing Scl1 level in plasma and *Streptococcus* level in stool samples before and after fecal microbiota transplantation (FMT) in SAH patients.

**Results:** Stool metaproteomics showed significant increase in 10 bacterial genera (*Streptococcus, Staphylococcus,* and *Clostridium*) in SAH→Healthy-rats, mirroring changes seen in ALD-rats (FC>1.5, p<0.05). Clusters of Orthologous Groups analysis indicated increased post-translational modifications (PTMs) and decreased lipid metabolism in ALD-rats and SAH→Healthy-rats. Liver kinome profiling showed 85 upregulated kinases in SAH→Healthy-rats, with 34 overlapping with ALD-rats, associated with inflammation (Mapk14, Map3k10 and others), fibrosis (Tgfbr1, Igf1r, and others), lipid metabolism (Cdk14, Cdk18) and regeneration (Met, Bmpr1a and others). FMT from healthy human-donors to ALD-rats reversed the expression of *Streptococcus* (10-fold), *Staphylococcus* (3-fold), and *Clostridium* (5-fold), along with 18 kinases associated with inflammation (Btk, Camk4, Nuak1) and fibrosis (Tgfbr1, Col4a1, Fgfr2 and others). Strong association (r²>0.9, p<0.05) was seen between *Streptococcus* abundance and fibrosis and inflammation-related kinases. Level of Scl1 was significantly high in SAH and ALD-rat stool samples (FC>2, p<0.05). It showed strong affinity for TGFβR1, as validated by molecular docking (>86%confidence) and immunoprecipitation assays (>100FC,p<0.05), indicating it as a potential ligand for TGFβR1. Concordantly, expression of Scl1 was highest amongst all the known ligands for TGFβR1 (p<0.05) suggesting Scl1 is a major contributor for TGFβR1 activation and its downstream signalling in these patients. Following fecal microbiota transplantation, expression of Scl1 in plasma and *Streptococcus* levels in stool were significantly reduced by 2.1-folds and 1.9 folds respectively in SAH patients. Further, Scl1 levels SAH plasma is capable of predicting poor therapeutic response and 30-day mortality with high accuracy (AUC=0.92, cutoff >70 normalized abundance).

**Conclusion:** Alcohol induced gut dysbiosis alters the liver kinome, promoting inflammation, fibrosis, and lipid dysregulation. Streptococcal Scl1 is identified as a bacterial protein mimicking human collagen and capable of activating fibrotic signalling pathway through TGFβR1in liver and is capable of predicting poor response in SAH. FMT from healthy donor restores microbial imbalance and harmful microbial-host interactions.

## Introduction

Alcohol-associated liver disease (ALD) is a progressive disorder encompassing steatosis, alcohol associated hepatitis (AH), severe AH (SAH), fibrosis, cirrhosis, and acute-on-chronic liver failure (ACLF) ^1^. While excessive alcohol consumption is the primary driver, growing evidence highlights the pivotal role of gut dysbiosis in its pathogenesis by increasing intestinal permeability and facilitating the translocation of microbial products such as lipopolysaccharides (LPS) into the portal circulation ^2^. This process triggers hepatic immune activation, inflammation, and fibrosis, accelerating disease progression^2^.

Gut microbiota, comprising of bacteria, fungi, archaea, and viruses, maintains intestinal homeostasis and regulates hepatic metabolism^3^. Alcohol consumption skews microbial composition, promoting the overgrowth of pathogenic bacteria such as *Enterobacteriaceae, Proteobacteria*, and *Veillonella* while depleting beneficial species like *Lactobacillus* and *Faecalibacterium prausnitzii*^4,5^. Notably, *Escherichia coli* and *Klebsiella pneumoniae* are linked to poor prognosis in severe ALD, exacerbating systemic endotoxemia and immune dysregulation^6^.

Preclinical studies confirm that transferring gut microbiota from alcohol-fed mice and or microbiota from SAH patients to germ-free mice (referred as humanised mice) induces hepatic inflammation and fibrosis, reinforcing the role of gut bacteria in ALD^7,8^. Beyond inflammation, gut microbiota profoundly influence hepatic fibrosis, a key driver of ALD progression^9^. Certain pathogenic bacteria, such as S*treptococcus pneumoniae* and *Streptococcus pyogenes*, have been implicated in pulmonary fibrotic diseases through their activation of the transforming growth factor-beta (TGFβ) signalling pathway^10,11^. Though the exact mechanism associated with TGF-beta activation and induction of fibrosis in alcohol associated liver diseases is obscure and warrants further analysis.

Kinases regulate cellular functions via phosphorylation and play crucial roles in the pathogenesis of ALD^12,13^..

We hypothesized that that alcohol-associated gut microbiota (AGM) may impact kinome signalling in the liver to drive inflammation and fibrosis. Using metaproteomics analysis in the stool sample and kinome analysis in the liver we identified that AGM, particularly *Streptococcus* species, secrete Scl1, a collagen-like protein which induces fibrosis by binding to TGFβ receptor 1 (TGFβR1). Scl1 activates p38 MAPK pathway and induces hepatic inflammation and fibrosis. Fecal microbiota transplantation (FMT) was found to mitigate these ill-effects.

These novel findings highlight the interplay between the microbial proteins released by alcohol ingestion and their effect on and liver kinases in development of liver disease. This offers a new perspective on the pathogenesis of ALD. Targeting these kinases dysregulated by AGM could serve as a novel therapeutic strategy to mitigate inflammation and fibrosis in ALD.

## Methods

### Ethics

All animal procedures were approved by the Institutional Animal Ethics Committee (IAEC), with the approval code IAEC/ILBS/007 for the ethanol administration to rats. Human samples enrolled in the study were approved by Institutional Review Board-IEC/2022/93/MA08 and IEC/2022/93/MA09.

Data regarding FMT in SAH patients was taken from previously published study Apoorva et. al. from our centre. 16s metagenomics and proteomics data (pre-and post-FMT samples) were re-analysed for the expression of *Streptococcus* bacteria in the stool samples (n=16) and Scl1 protein in plasma.

### Study Design

This study aimed to explore the impact of AGMs on liver kinase signalling. This was achieved by employing comprehensive experimental design integrating stool metaproteomics and liver kinome profiling in the following study groups: i) ALD rats, ii) healthy rats gavaged with stool slurry of SAH patients referred as SAH→healthy-rats (HR), iii) ALD rats gavaged with stool slurry of healthy human-donors referred as Human donor→ALD-rats.

Kinases activated by AGMs were further validated using molecular docking analysis and functionally assessed by treating THP-1 monocytes with bacterial secretome, followed by co-immunoprecipitation coupled with mass spectrometry (Co-IP MS). Additionally, stool of SAH patients (n=24) and liver biopsy (n=19) samples were analysed for the expression of identified bacterial proteins interacting with liver kinases. The therapeutic potential of FMT was also investigated by evaluating changes in bacterial abundance and corresponding protein secreted by bacteria in pre- and post-FMT stool and plasma samples of 16 SAH patients, using 16s metagenomic and proteomic approaches. This data was re-analysed from previously published datasets generated at our center ^14^ (Figure1A).

### Severe alcohol-associated hepatitis patient selection criteria

Patients with SAH were included based on: (1) history of chronic alcohol use, (2) recent onset of jaundice, (3) Maddrey’s Discriminant Function (DF) >32, and (4) serum bilirubin >3 mg/dL and AST/ALT >3 times ULN. Patients with active infections, hepatocellular carcinoma, or other chronic liver diseases were excluded. Following things were analysed in SAH samples: Scl1 protein level in SAH stool (n=24), Scl1 protein level in SAH liver biopsy (n=19) procured from National liver disease biobank, New Delhi. *Streptococcus* levels in SAH stool samples (n=10), Scl1 and *Streptococcus* levels in pre-and post-28-day FMT plasma and stool samples of SAH patients (n=16) respectively. This data was re-analysed from the previously published data from our study^14^. Scl1 levels in plasma of SAH were further validated in another cohort of 88 patients (table S5).

FMT donors were selected based on previously defined criteria^15,16^.

### Animals

In this study, 5-6□weeks-old male and female Long Evans rats, weighing 200-250□g, were obtained from Animal Facility of the National Brain Research Centre, India and inhabited in animal facility at ILBS. All rats were housed in a room under a 12□h regular light/dark cycle at an ambient temperature 21°C. The animal studies were approved by the Institutional Ethics Committee of ILBS and conducted in accordance with ethical standards.

### Establishment of chronic alcohol-related liver disease rat model (ALD)

Following a one-week acclimatization period during which rats were provided with water and food ad libitum, they were transitioned from a standard chow diet to the Lieber-De Carli (LDC) control liquid diet (Catalog# F1259, BioServ, Frenchtown, NJ). This transition was conducted in accordance with guidelines set by the National Institute on Alcohol Abuse and Alcoholism (NIAAA) ^17^. After the transition, the rats were randomly assigned to two groups: a control group (n=6) and an ALD group (n=6). The ALD group underwent a gradual introduction to ethanol over a seven-day period, starting with a concentration of 5% and progressively increasing to 37%. This ethanol regimen was maintained for 24 weeks. Meanwhile, the control group was pair-fed with the LDC control diet, ensuring that their caloric intake matched that of the ALD group, with non-alcoholic components substituted for the ethanol-derived calories.

### Reciprocal Fecal Microbiota Transplant: Animal study

Reciprocal Fecal Microbiota Transplant referred to as a bidirectional FMT study. The main aim of this study was to determine how different microbiota compositions (e.g., SAH vs. healthy donors) influence host physiology, such as liver inflammation or fibrosis.

The rats were randomly assigned to one of three groups, with 6 rats in each group:

1. Human donor→Healthy-rats: Healthy rats that served as a control vehicle group, receiving stool slurry from a healthy human donor.
2. SAH→Healthy-rats: Healthy rats that received stool slurry from patients with SAH (Demographics in Table S4).
3. Human donor→ALD-rats: Rats with alcohol-related liver disease (ALD-rats) that received stool slurry from a healthy human donor.

Fecal microbiota transplantation (FMT) was performed daily for one week. After the completion of the one-week FMT regimen, the rats were sacrificed for further analysis.

### SAH patient stool sample collection and preparation

The stool of SAH patients and healthy donor subjects were collected and stool slurry was prepared in normal saline under aseptic conditions. Aliquots of 100mg/ml were made and were stored in −80^0^ until use.

### Fecal microbiota transplantation protocol

Rats were kept on fasting overnight before performing bowel wash. Bowel wash was performed 4 times with 1mL of 425g/l PEG at interval of 20 minutes by oral gavage. 1mL of 100mg/mL stool was dissolved in normal saline and was administered to rats by oral gavage every day for 1 week.

### Stool sampling

Stool and fecal samples from patients and rats respectively were collected and immediately lysed in lysis buffer to perform targeted proteomic and metaproteome profiling. Remaining fecal samples were stored in liquid nitrogen.

Liver tissue and plasma samples of rats and liver biopsies from patients were collected and stored in liquid nitrogen until use.

### Metaproteome profile of Fecal and stool samples

Metaproteome profiling was conducted on fecal samples collected from the control rats, ALD-rats, SAH→Healthy-rats, and human donor→ALD-rats and SAH patients (stool and liver). The fecal samples were dissolved in lysis buffer and subjected to sonication (3 cycles of 10 seconds on, 10 seconds off, at an amplitude of 30). The resulting supernatant was collected, and 50 µg of protein was measured for further processing. The protein samples were first reduced with 10 mM DTT at 60°C for 1 hour, followed by alkylation with 10 mM iodoacetamide (IAA) in the dark at room temperature. The samples were then digested with trypsin (Promega: V5280) for 24 hours at 37°C.

After digestion, the samples were desalted using C18 spin columns (PierceTM: 89870) and lyophilized. The lyophilized peptides were reconstituted in 0.1% formic acid and then subjected to nano-electrospray ionization followed by tandem mass spectrometry (MS/MS) using a Q-ExactiveTM Plus instrument (Thermo Fisher Scientific, San Jose, CA, United States). Peptides were initially enriched on a trap column (75 µm x 2 cm, 3 µm, 100Å, nano Viper 2Pk C18 Acclaim PepMapTM 100) at a flow rate of 8 µl/min. They were subsequently separated on an analytical column (75 µm × 25 cm, 2 µm, 100Å, nano Viper C18, Acclaim PepMapTM RSLC). Peptides were eluted over a 120-minute gradient (3–95% buffer B: 80% acetonitrile in 0.1% formic acid) at a continuous flow rate of 300 nL/min. Mass spectrometry was performed using collision-induced dissociation with an electrospray voltage of 2.3 kV. Orbitrap analysis included full-scan MS spectra at a resolution of 70,000 over the m/z range 200–2000. Protein identification was carried out using the Mascot algorithm (Mascot 2.4, Matrix Science), with statistical significance determined at p<0.05 and q values (false discovery rate) also set at p<0.05, ensuring a false discovery rate threshold of 0.01^18^.

### Metaproteome data analysis

The MS/MS data were acquired and analysed by Proteome Discoverer (version 2.3, Thermo Fisher Scientific, Waltham, MA, United States) using the bacterial sequence (UniprotSwP_20170609 with sequences 467231 and MG_BG_UPSP with sequences 2019194). This was cross-validated using the Mascot algorithm (Mascot 2.4, Matrix Science) specifically for all possible microbial species. Significant peptide groups were identified at (p<0.05) and q-values (p<0.05), and the false discovery rate was at 0.01. Only rank-1 peptides with Peptides Sequence Match (PSMs)>3 were subjected to biodiversity and functional analysis using Unipept. Peptides mapping to the eukaryotic, fungal, and viral database were rejected and only bacterial species-associated peptides were segregated and subjected to statistical, functional, and biodiversity analysis as detailed in our recent publication ^18^. Identified metaproteins and corresponding list of bacteria is highlighted in Table S1 and S2 in respective study groups.

### Kinome profile in liver sample

Kinome profiling was performed on liver tissue samples from ALD rats, SAH→H rats, HD→H rats, and HD→ALD rats. The liver tissues were first pulverized using liquid nitrogen, and the resulting powder was dissolved in a lysis buffer containing a cocktail of protease (11836153001, Roche, cOmplete™, Mini Protease Inhibitor Cocktail) and phosphatase inhibitors. After removing contaminants, 1 mg of equivalent protein was taken for further analysis.

The protein samples were processed following the same digestion protocol as described in the metaproteome sample preparation. After digestion, the samples were desalted using C18 spin columns (PierceTM: 89852) and lyophilized. Phosphopeptides were enriched from the lyophilized samples using a combination of TiO2 (High selectTM TiO2 #A32993) and Fe-NTA phosphopeptide enrichment kit (High selectTM Fe-NTA #A32992) ^19,20^.

The enriched phosphopeptides were subjected to nano-electrospray ionization followed by tandem mass spectrometry (MS/MS) using a Q-ExactiveTM Plus instrument (Thermo Fisher Scientific, San Jose, CA, United States). Peptides were first enriched on a trap column (75 µm x 2 cm, 3 µm, 100Å, nano Viper 2Pk C18 Acclaim PepMapTM 100) at a flow rate of 8 µl/min and then separated on an analytical column (75 µm × 25 cm, 2 µm, 100Å, nano Viper C18, Acclaim PepMapTM RSLC).

The peptides were eluted over a 120-minute gradient (3–95% buffer B: 80% acetonitrile in 0.1% formic acid) at a continuous flow rate of 300 nL/min. Mass spectrometry was conducted using collision-induced dissociation with an electrospray voltage of 2.3 kV. The Orbitrap analysis included full-scan MS spectra at a resolution of 70,000, covering the m/z range from 200 to 2000

### Kinome data analysis

The MS/MS data were acquired and analysed by Proteome Discoverer (version 2.3, Thermo Fisher Scientific, Waltham, MA, United States) using Human sequence database. Mascot and Amanda with ptm-RS pipelines were used to identify the peptide sequences and corresponding proteins. Significant peptide groups were identified at (p<0.05) and q-values (p<0.05), and the false discovery rate was at 0.01. Only rank-1 peptides with Peptides Sequence Match (PSMs)>3 was considered. List of all the kinases were downloaded from Coral (http://phanstiel-lab.med.unc.edu/CORAL/) and were traced in our data. List of identified kinases in data set is enlisted in Table S3.

### Liver tissue histology

Liver tissue was processed and stained with hematoxylin-eosin (HE), Sirius red (SR), and Oil Red (OR) for histological examination following standard protocols.

### Statistical analysis

Statistical analyses were conducted using MetaboAnalyst 6.0 and GraphPad Prism-6. For comparisons between the two groups, unpaired (2-tailed) Student t-tests and Mann-Whitney U tests were employed. Correlations were assessed using Spearman correlation analysis, with coefficients (R²) greater than 0.9 considered significant. Features with p<0.05 are considered significant. Metaproteome data was analysed using Microbiomeanalyst 2.0 (https://www.microbiomeanalyst.ca/). For pathway analysis were performed using Enrichr, and Shinygo. Correlation analysis was performed using metaboanalyst and Metscape.

### Streptococcus secretome screening

We discovered Scl1 through a systematic approach. Streptococcus was found to be highly abundant in stool samples of SAH patients and ALD-rats, and its abundance strongly correlated with hepatic TGFβR1 expression. To investigate whether Streptococcus can activate TGFβR1, we cultured the bacteria and performed proteomic analysis of its secretome. Among the identified proteins, several showed sequence similarity to collagen. Literature mining further revealed that *Streptococcus* secretes a collagen-like protein, Scl1, known to mimic host collagen and could therefore potentially engage TGFβR1. This led to hypothesize and validate Scl1 as a key microbial factor capable of inducing TGFβR1 signalling.

### Sequence BLAST Analysis on NCBI BLAST for Scl1 and Collagen

The BLASTp search was performed on the NCBI BLAST server (https://blast.ncbi.nlm.nih.gov/Blast.cgi?PAGE=Proteins) to compare the Scl1 protein from *Streptococcus* with collagen sequence (Accession number: P02452). The FASTA sequences of both proteins were retrieved from UniProt and NCBI databases and saved for input. The default parameters were used, including the BLOSUM62 matrix, an E-value cutoff of 0.01, and standard gap penalties (−11, −1). The search was executed, and the results were analyzed based on sequence identity, query coverage, and E-values. Significant hits aligning with collagen-like sequences were identified. Alignments were examined for conserved Gly-X-Y motifs, characteristic of collagen, to confirm potential mimicry by Scl1.

### Molecular Docking of TGFβR1 and Scl1 Protein Using HDOCK Server

Molecular docking of the TGFβR1 receptor and Scl1 protein was performed using the HDOCK server to analyze their binding interactions. The HDOCK server was accessed via http://hdock.phys.hust.edu.cn/, and the TGFβR1 receptor and Scl1 ligand were uploaded in FASTA format. The protein-protein docking mode was selected, and the docking simulation was performed using the default global docking parameters, allowing free docking without predefined binding constraints. The sequence-based template search was enabled for better accuracy. The job was submitted, and the docking simulation was monitored until completion.

Upon completion, the top-ranked docking models were analyzed based on their confidence scores, RMSD values, and binding energy. The highest-ranked docking pose was selected, considering the lowest binding energy and highest confidence score.

The docking results provided insights into the binding site and interaction pattern of Scl1 with TGFβR1, supporting the hypothesis that Scl1 may modulate TGFβ signaling. The results were further validated through an in vivo experiment, where the concentrated bacterial secretome was co-cultured with the THP-1 monocyte cell line. Following this, TGFβR1 was immunoprecipitated, and the pulled-down lysate was analyzed using mass spectrometry to detect the presence of Scl1 protein. The sequence coverage of Scl1 was then traced to confirm its interaction with TGFβR1.

### Streptococcus pneumonia^21,22^ and S. pyogenes^22^ culture and secretome collection

The *Streptococcus pneumoniae* wildtype strain D39 (NCTC 7466; serotype 2) was obtained from the National Type Culture Collection (NTCC), United Kingdom. Secretome preparation was carried out with minor modifications to the protocol described by Jhelum et al □1]. Briefly, a primary culture was initiated in Todd-Hewitt broth supplemented with 0.5% yeast extract (THY) using glycerol stocks stored at –80°C. Cultures were incubated at 37°C in a 5% CO□ atmosphere until they reached mid-logarithmic phase (OD□□□ = 0.4–0.6). Subsequently, a secondary culture was established by inoculating fresh THY medium with 1% (v/v) of the primary culture. In parallel, uninoculated THY medium was incubated under identical conditions to serve as a negative control. Cultures were allowed to grow until an OD□□□ of 0.6 was reached, at which point bacterial cells were pelleted by centrifugation at 13,000 × g for 10 min at 4°C. The supernatants from both the bacterial culture and the negative control were carefully collected, avoiding disturbance of the pellet and subsequently filtered through 0.2□µm sterile syringe filters to eliminate any remaining bacterial cells. To confirm the sterility of the secretome preparations, aliquots of the filtered supernatants were plated on tryptic soy agar plates supplemented with 5% (v/v) defibrinated sheep blood and incubated overnight at 37°C in a 5% CO□ atmosphere. Only sterile, contamination-free preparations were used in downstream analyses.

*S. pyogenes* was obtained from MTCC, strain 14289 was cultured using same protocol as mentioned above for *S. pneumonia*.

Supernatant was filtered through 0.2µ filter to avoid any bacterial contamination. Secretome as well as THY control were plated on Tryptic soy agar (TSA) plates with 5% sheep blood to negate any bacterial contamination.

Plates images are shown below observed after (A) 24 H and (B) 60 H of plating *Streptococcus pneumonia* and *Streptococcus pyogenes*.

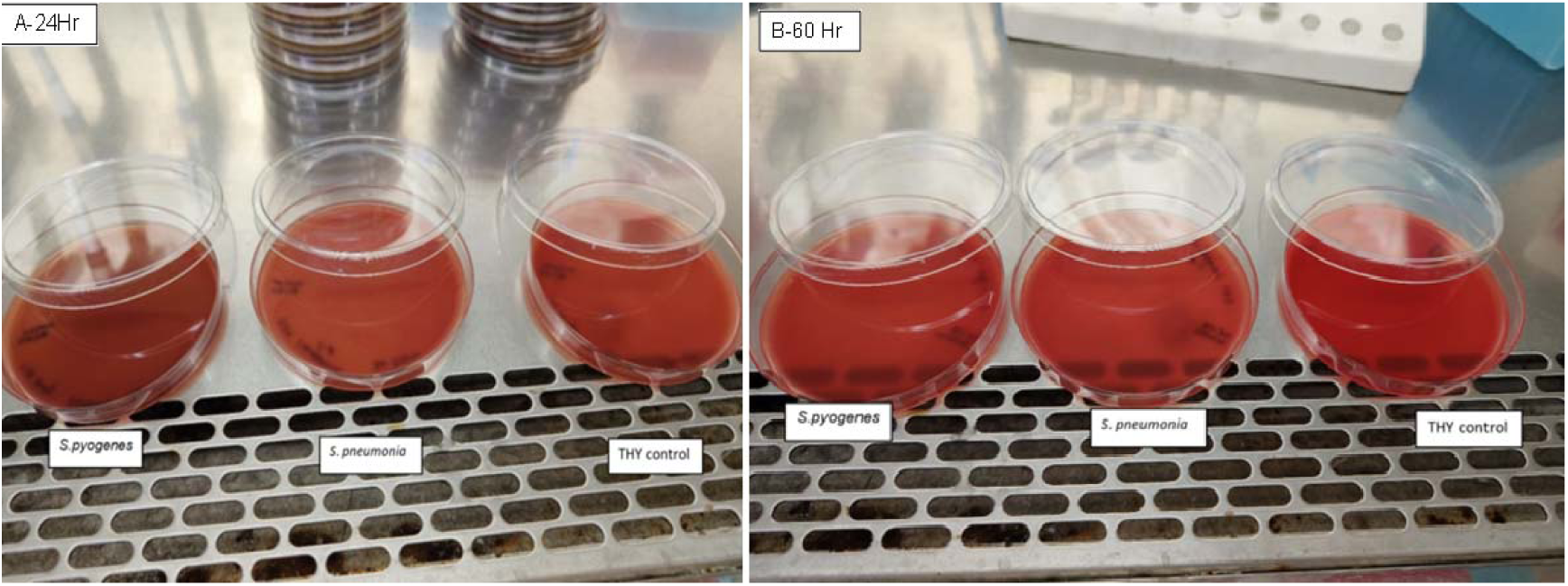

### THP1 cell line culture

#### Treatment of Activated THP-1 Macrophages with Bacterial Secretome

THP-1 cells were cultured and activated with 10 ng/mL PMA and incubated for 24 hours, allowing differentiation into macrophages. The bacterial secretome from *S. pneumoniae* and *S. pyogenes* was collected and concentrated from 10 mL to 1 mL using 10kDa amicon filter. The activated THP-1 macrophages were then treated with 10% (v/v) of the concentrated bacterial secretome and incubated for an additional 24 hours. Following treatment, the cells were lysed using NTN buffer (100 mM NaCl, 20 mM Tris, pH 7.4, 0.5% NP-40, 10% glycerol, 1X PMSF, and 1X protease inhibitor cocktail). The lysate was then subjected to immunoprecipitation using anti-TGFβR1 antibodies, followed by further analysis using mass spectrometry technique.

#### Immunoprecipitation (IP) Protocol

The immunoprecipitation of TGFβR1 was performed using agarose beads and anti-TGFBR1 antibody (PAA397Hu01, Cloud clone) to isolate the target protein. The experiment was carried out under cold conditions to preserve protein integrity.

##### Preclearing of Lysate

∼1mg protein was taken and 30 µL of protein A agarose beads (SC-2001) was added, followed by 500 µL of NTN buffer (without NP-40). The tubes were centrifuged at 500 g for 2 minutes at 4°C, and the supernatant was discarded. The prepared lysate was then added to the beads and incubated on rotator for 1hr followed by centrifugation at 12000g for 5 min. Beads were discarded and anti-TGFBR1 antibody was added (3ug) and incubated overnight on rotator, followed by washing with NP40 buffer at 5000rpm.

##### Elution for Intact Protein Recovery

To elute the bound proteins, the beads were resuspended in 50–100 µL of 50 mM ammonium bicarbonate (pH 8.0). The samples were incubated at room temperature for 10–15 minutes with gentle shaking. The tubes were centrifuged, and the supernatant containing the eluted proteins was collected. The collected samples were processed for mass spectrometry analysis.

#### Targeted Proteomics

Stool and liver samples from rat models and SAH patients were processed for targeted proteomics analysis. The protein sequences of Scl1 were retrieved in FASTA format and provided to Protein Discoverer for peptide identification. The identified peptides were mapped onto the Scl1 protein sequence to confirm its presence in stool and liver samples, and the peptide spectral matches (PSMs) were analyzed to determine relative protein abundance. The results were statistically evaluated to assess differential expression between control and diseased conditions^18^.

## Results

### Development of ALD-rat model and Reciprocal Fecal Microbiota Transplant Design workflow

This study aimed to examine how AGMs impacts the liver kinome and contribute to inflammation and fibrosis, and whether FMT can reverse these effects. This was achieved by performing reciprocal FMT (Figure 1A). We analysed stool metaproteome and liver kinome across ALD-rats, SAH→Healthy-rats (healthy rats gavaged with stool from SAH patients), and human donor→ALD-rats (ALD rats receiving stool from healthy human donors) to identify microbiota-driven kinase alterations (Figure 1A).

**Figure 1:**
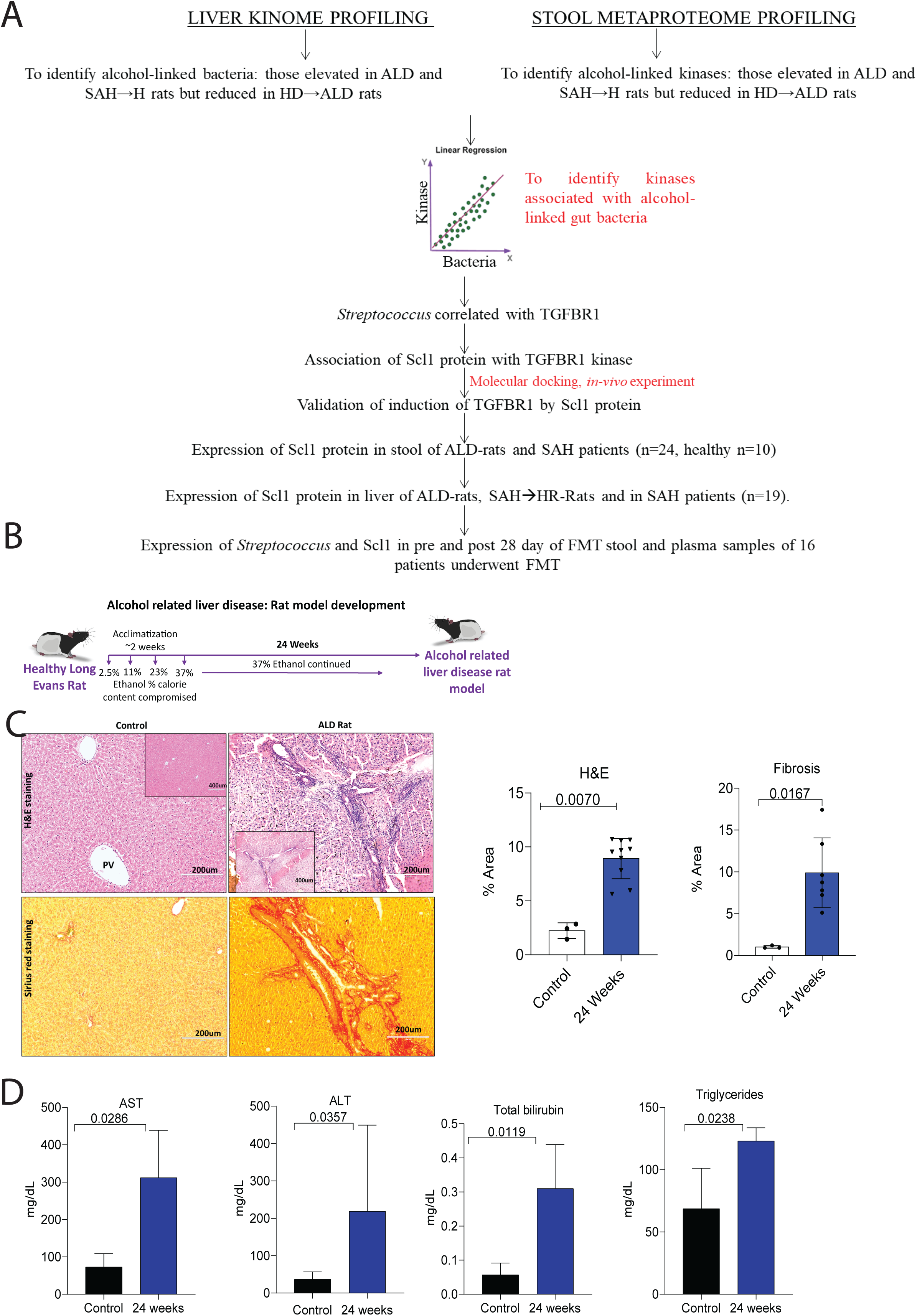
Development of ALD model and Criss cross FMT workflow. 1A. Diagrammatic representation of study design. 1B. Diagrammatic representation of ALD rat model development 1C. Histological assessment of liver injury and fibrosis in ALD rats at 6 months. 1D. Biochemical evaluation of liver function and lipid metabolism in ALD rats at 6 months.

We integrated stool metaproteome and liver kinome data to identify microbiota-regulated kinases. Bacterial secretome proteomics, molecular docking, Co-IP-MS/MS, and THP-1 cell assays and validation in human FMT patients revealed microbial proteins induces kinase. This integrative approach uncovers a mechanistic link between gut microbiota, kinase dysregulation, and fibrotic signalling in alcohol-related liver disease (Figure 1A).

Development of ALD-rat model involved chronic administering of 37% ethanol diet for six months (Figure 1B). Histological analysis revealed a significant increase in the percentage of inflammation and fibrosis (Figure 1C). Additionally, biochemical parameters, including AST, ALT, total bilirubin, and triglycerides, were significantly elevated, confirming that this model effectively mimics ALD and is suitable for studying alcohol-associated liver disease (Figure 1D).

### Comparative metaproteome and functional analysis in stool of ALD-rats and humanized SAH→Healthy-rats

This study used a comprehensive approach to identify microbial signatures linked to alcohol-associated gut dysbiosis (Figure 2A). As compared to controls rats, the stool metaproteome analysis of ALD-rats and SAH→Healthy-rats revealed similar microbial profiles, with increased *Bacillota* and *Bacteroidota* and decreased *Actinomycetota* (p<0.05, Figure 2B). Shannon beta diversity showed overlapping microbiota at genus level between ALD-rats and SAH→Healthy-rats, indicating shared composition. Both groups also exhibited significantly reduced alpha diversity^23^ (p<0.05), confirming SAH→Healthy-rats as an effective model for studying ALD-associated gut dysbiosis (Figure 2C). Taxonomic analysis showed significant upregulation of *Campylobacterota, Pseudomonadota, Bacillota, Actinomycetota,* and *Bacteroidota* in stool of both ALD-rats and SAH→Healthy-rats (Figure 2D). Linear Discriminant Analysis (LDA) identified *Campylobacterota, Pseudomonadota, Bacillota, Actinomycetota,* and *Bacteroidota* as the most discriminative phyla in both the groups compared to control (Figure 2E). At the genus level, *Streptococcus*, *Clostridium, Corynebacterium, Mycobacterium,* and *Staphylococcus* were significantly enriched in stool of both the groups (p<0.05, Figure 2F), suggesting their potential role in alcohol-associated gut dysbiosis. COG analysis showed increased activity in pathways related to metabolism, PTMs, and chaperones (Figure 2G). These findings highlight the SAH→Healthy-rats model as a robust tool to study gut microbiota–driven mechanisms in ALD pathogenesis and therapeutic targeting.

**Figure 2:**
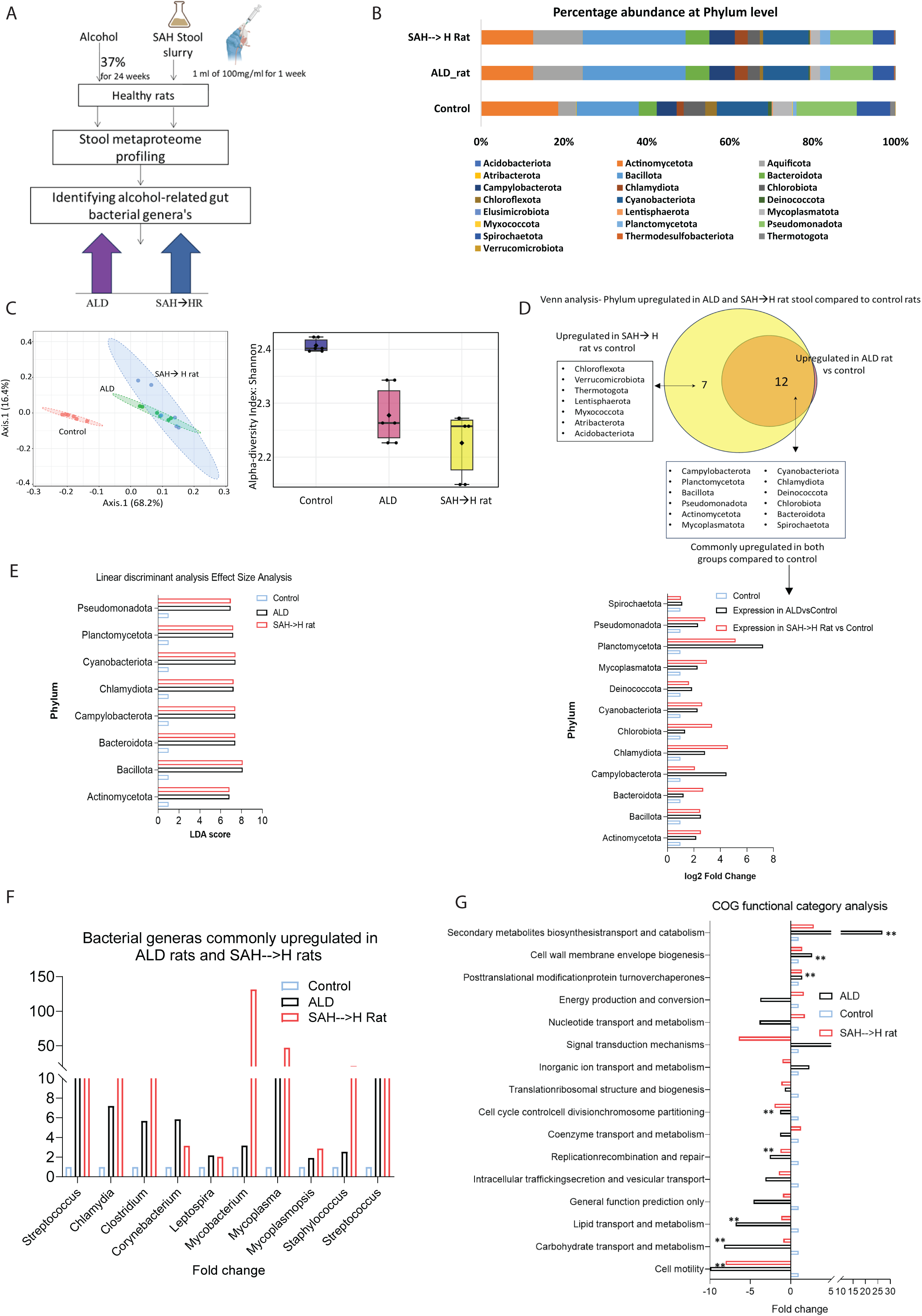
Comparative metaproteome and functional analysis of ALD Rats and humanized SAH→H Rats. 2A. Schematic representation of the experimental approach to identify alcohol-associated gut microbiota in ALD. 2B. Stool metaproteome composition at the phylum level reveals similar phylum abundance in ALD and SA→HR rat stool. 2C. Overlapping beta diversity and significantly reduced alpha diversity in ALD and SA→HR rat microbiota indicating a shared dysbiotic microbial community structure between the two groups. 2D. Venn analysis and bar plot illustrating the bacterial phyla commonly upregulated in ALD rat stool and SA→HR rat stool samples (p<0.05, FC>1.5 compared to control). 2E. Linear Discriminant Analysis (LDA) scores highlighting bacterial phyla similarly enriched in ALD and SA→HR rat stool samples. 2F. Bar plot showing fold change expression of bacterial genera commonly upregulated in ALD stool and SAH-->H rat stool (FC>1.5, p<0.05). 2G. COG functional classification of stool metaproteins reveals activation of similar microbial pathways in ALD and SAH→HR rats.

### Comparative Liver kinome Profiling and functional analysis of ALD-rats and humanized SAH→Healthy-rats

This study used systems-level kinome profiling approach to examine signalling changes driven by AGMs in liver (Figure 3A). Principal component analysis (PCA) revealed a distinct clustering of kinome profiles in both ALD-rats and SAH→Healthy-rats, indicating a global shift in kinase activity associated with disease state (Figure 3B). ALD-rats showed 172 upregulated and 97 downregulated kinases while SAH→Healthy-rats showed 85-upregulated and 69-downregulated kinases (FC>1.5, p<0.05, Figure 3C). ALD-associated kinases were associated with RTK, MAPK, TLR, and TGF-β signalling, while autophagy pathways and others were suppressed (Figure 3D). SAH→Healthy-rats showed enhanced immune and inflammatory signalling (NF-κB, TCR, IL-17) and reduced JAK-STAT and calcium signalling (Figure 3E). Thirty-four kinases overlapped across models, including TGFβR1, MAP2K1, IGF1R, and BTK key regulators of inflammation, fibrosis, and metabolism (Figure 3F). These findings reveal that AGMs drives hepatic kinase dysregulation linked to inflammation and fibrosis, highlighting potential therapeutic targets for ALD.

**Figure 3:**
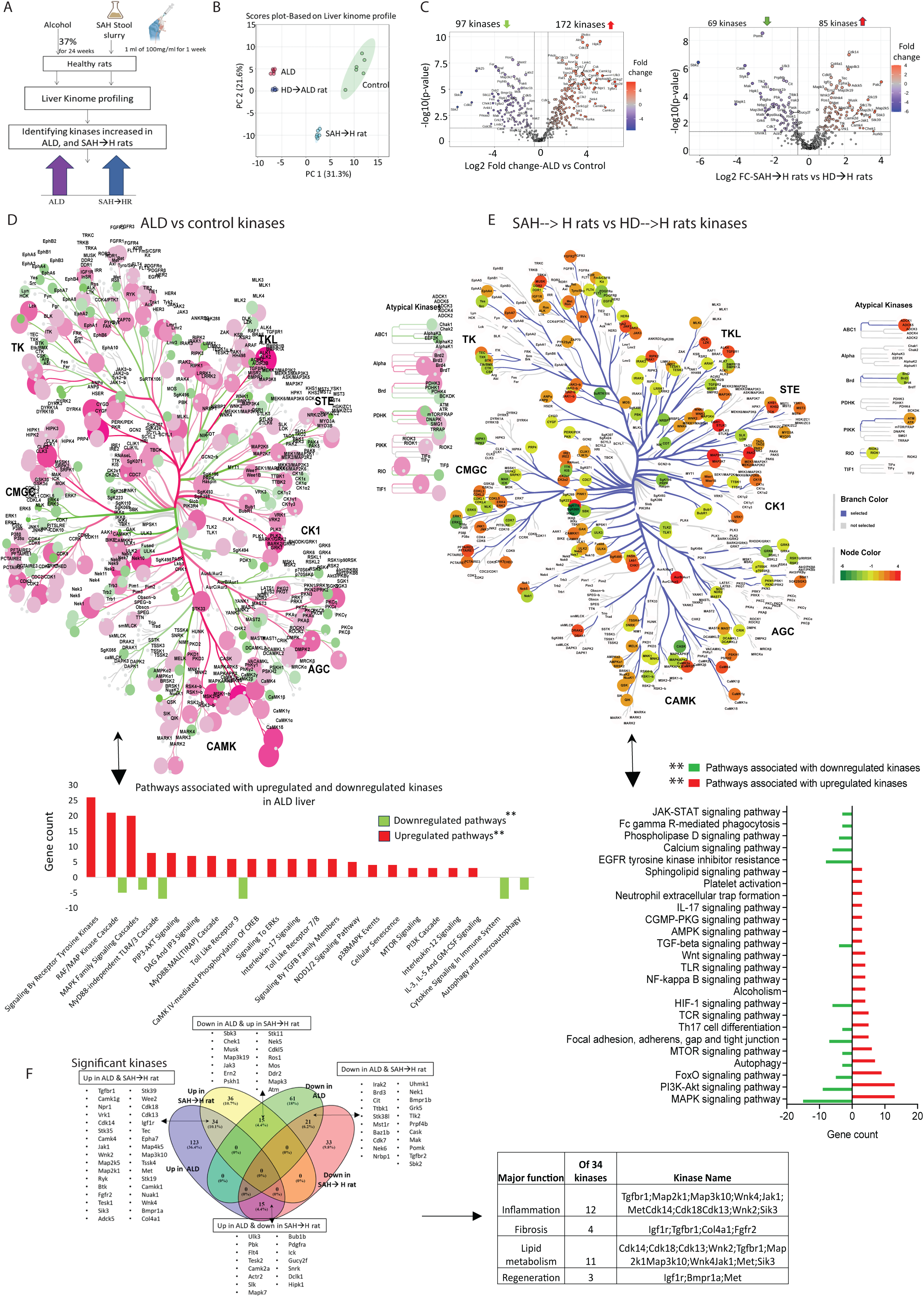
Comparative Liver kinome Profiling and functional analysis of ALD Rats and humanized SAH→H Rats model. 3A. Schematic representation of the experimental approach to identify kinases in liver linked with alcohol-associated gut microbiota in ALD. 3B. Principal Component Analysis of liver kinome profiles in Control, ALD, HD→ALD, and SAH→HR rats 3C. Volcano plot showing differentially expressed kinases in liver of ALD rats compared to control and SAH→H rats compared to HD→H rats. 3D. Kinome map highlighting differentially expressed kinases in ALD liver compared to control rat liver (pink=upregulated, green= downregulated, FC±1.5, p<0.05). Bar plot shows pathways associated with differentially upregulated and downregulated kinases. 3E. Kinome map highlighting differentially expressed kinases in SAH→H rats liver compared to HD→H rat liver (pink=upregulated, green= downregulated, FC±1.5, p<0.05). Bar plot shows pathways associated with differentially upregulated and downregulated kinases. 3F. Venn analysis showing 34 kinases commonly upregulated in ALD rat and SAH→H rat liver associated with inflammation, fibrosis, lipid metabolism and regeneration.

### Healthy FMT restores bacterial modulated liver kinome alteration in preclinical rat model of ALD

To validate the contribution of AGMs in modulating hepatic kinase activity, we performed FMT from healthy human donors to ALD-rats (human donor→ALD-rats Figure 4A). Of the 34 kinases upregulated in both ALD-rats and SAH→Healthy-rats, 18 kinases showed significant downregulation in the human donor stool→ALD-rats group. Notably, kinases such as BTK, CAMK4, MAP2K5, MAP4K5, IGF1R, and TGFβR1 all of which are implicated in inflammation, fibrosis, and metabolic dysregulation were also reversed (Figure 4B). These findings align with prior studies reporting the role of gut-derived microbial products in activating hepatic kinase pathways, including TGFβ signalling, during chronic liver injury^24^. A significant reversibility of taxa specific to SAH-stool exposure (*Streptococcus, Staphylococcus, Mycoplasma, Corynebacterium,* and *Clostridium*) was seen upon healthy FMT in human donor→ALD-rat stool (Figure 4C).

**Figure 4:**
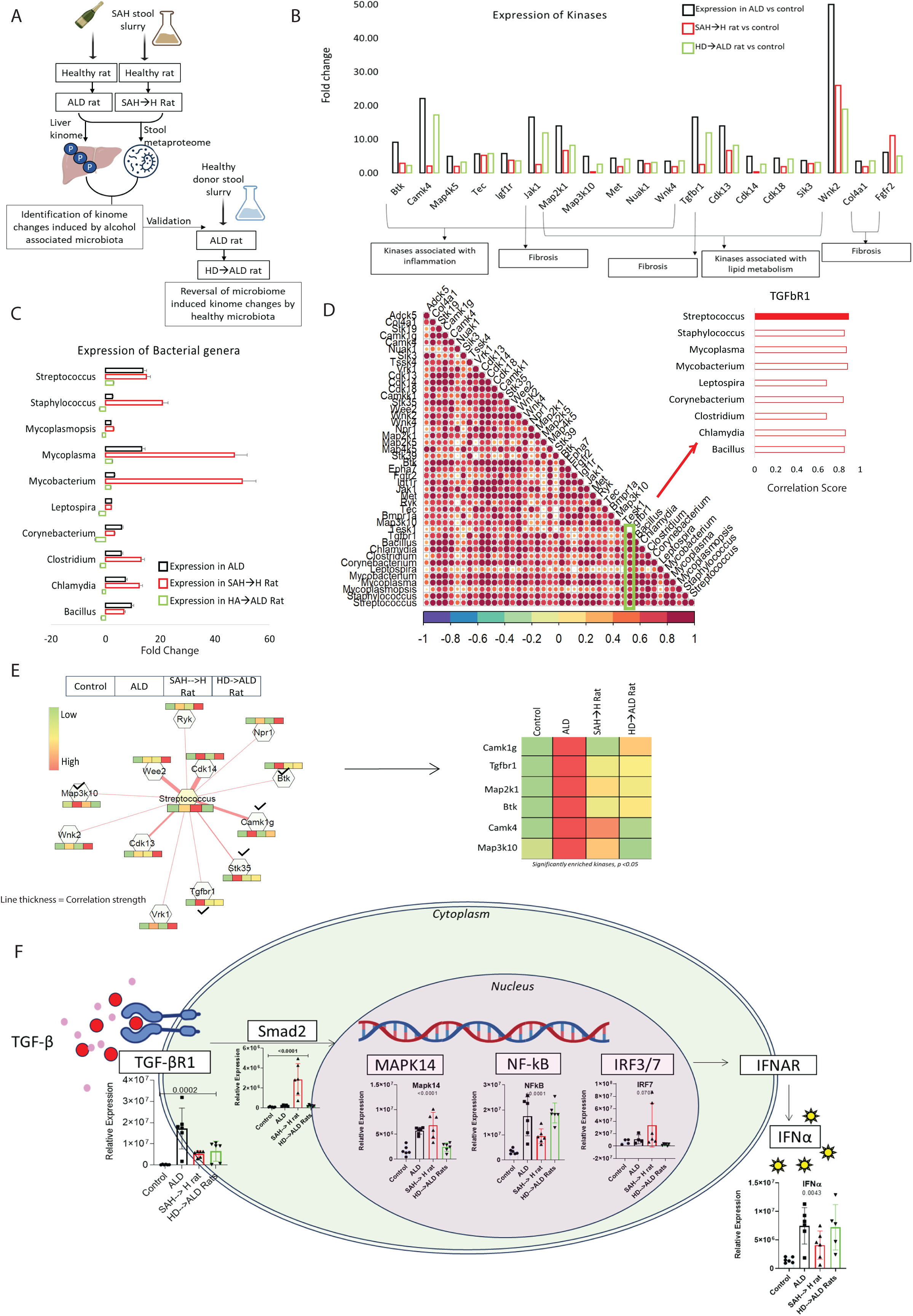
Healthy FMT restore bacterial modulated liver kinome alteration in preclinical model of ALD. 4A. Schematic presentation of approach used to identify the kinases in liver associated with alcohol associated gut microbiota. 4B. Bar plot showing expression of kinases commonly upregulated in ALD and SAH→H rat liver but got reversed in HD→ALD rat liver. 4C. Bar plot showing expression of bacterial genera commonly upregulated in ALD and SAH→H rat stool but got reversed in HD→ALD rat stool. 4D. Correlation plot showing correaltion between bacterial genera and kinases linked to alcohol associated gut microbiota. TGFBR1 showed correlation of R^2^=0.88 with *Streptococcus*. 4E. Correlation of Streptococcus with kinases linked to alcohol associated gut microbiota. Heat map shows expression of these kinases in control, ALD, SAH→H rats and HD→ALD rats liver. Only kinases with significant change were considered. 4F. Expression of proteins associated with TGFBR1 and inflammation axis.

Stool microbiota and liver kinome cross correlation analysis showed a strong positive association between *Streptococcus* and TGFβR1 (r=0.88; Figure 4D), suggesting that *Streptococcus* may play a central role in activating TGFβ signalling in the context of alcohol-associated dysbiosis. Additional inflammatory kinases such as BTK, MAP3K10, STK35, and CAMK1G also showed significant correlation with *Streptococcus* abundance (Figure 4E), indicating a broader role of this genus in driving inflammation.

Stool from SAH patients increased hepatic expression of TGFβR1 effectors (SMAD2, MAPK14, NFκB, IRF, IFN-α) in healthy rats in a similar trend seen in ALD-rats), and was reversed post FMT from healthy donor was given to ALD-rats (Figure 4F). These results suggest that alcohol-altered gut microbiota, particularly *Streptococcus*, drives TGFβR1-mediated inflammation and fibrosis. FMT-mediated restoration of microbial balance highlights the therapeutic promise of targeting gut-liver signalling in ALD.

### Identification and Validation of Scl1 as a Streptococcus-Derived Activator of TGFβR1 Signalling in Alcohol-Associated Liver Disease

To decipher the molecular mechanism by which *Streptococcus* contributes to the activation of hepatic TGFβR1 in ALD, we adopted a systematic approach outlined in Figure 5A.

**Figure 5:**
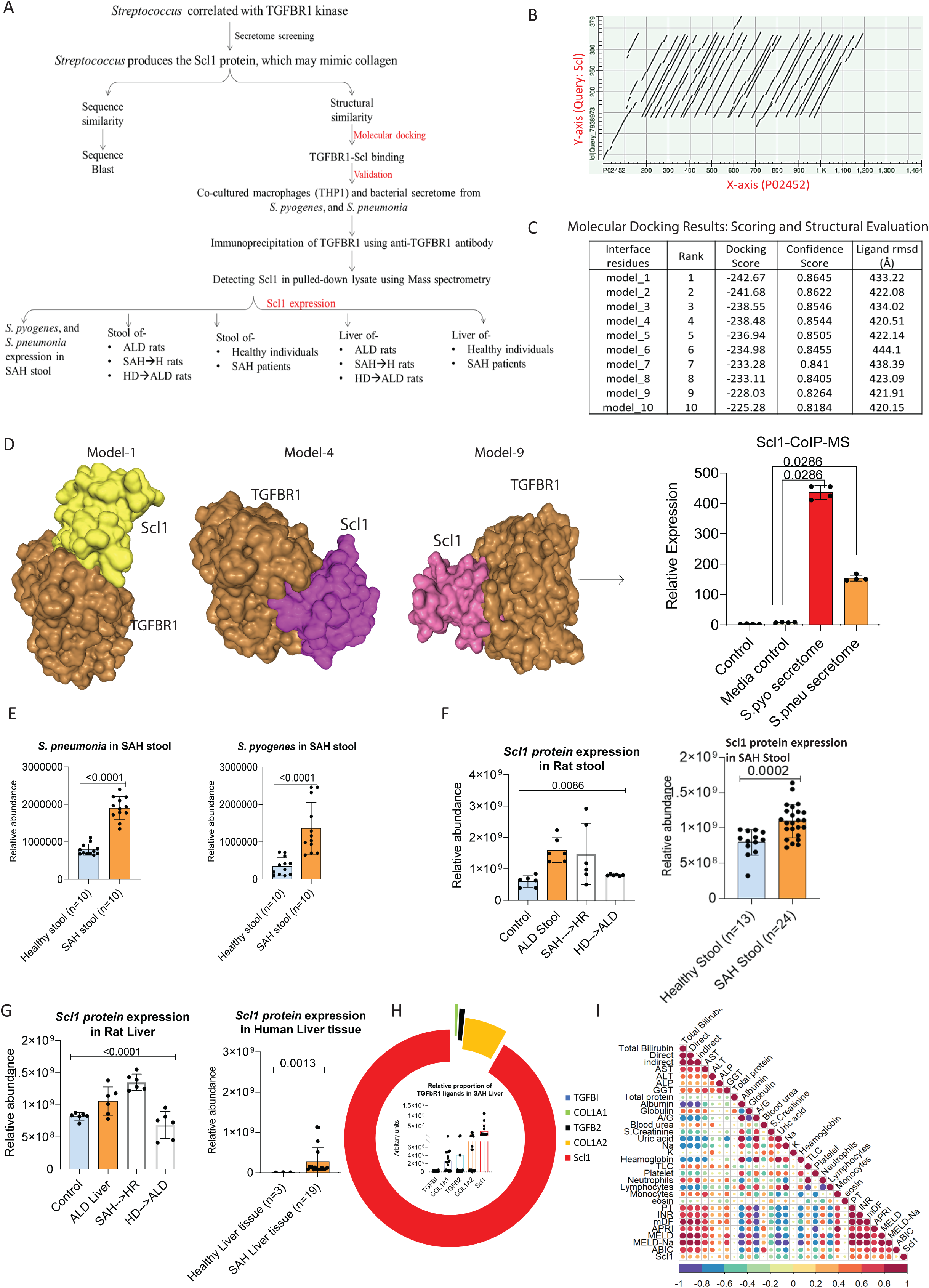
Identification and Validation of Scl1 as a Streptococcus-Derived Activator of TGFβR1 signalling in Alcohol-Associated Liver Disease: 5A. Flow diagram representing the approach used to show association between *Streptococcus* and TGFBR1. 5B. Dot plot showing sequence similarity between the query protein Scl (Y-axis) and human collagen alpha-1(I) chain (P02452) (X-axis). 5C. Table summarizing the top 10 predicted docking models of protein-ligand complexes ranked based on docking score. 5D. Docked structure illustrating the interaction between TGFβR1 and Scl1 protein. The accompanying bar plot quantifies Scl1 enrichment in a co-immunoprecipitation followed by mass spectrometry (Co-IP–MS) assay using anti-TGFβR1 antibody. 5E. Expression of *S. pneumonia* and *S. pyogenes* in stool samples of healthy individuals (n=10) and SAH patients (n=10). 5F. Expression of Scl1 protein in stool samples of control, ALD rats, SAH→H rats, HD→ ALD rats and stool samples of healthy individuals (n=13) and SAH patients (n=24). 5G. Expression of Scl1 protein in liver samples of control, ALD rats, SAH→H rats, HD→ ALD rats and liver tissue of healthy individuals (n=3) and SAH patients (n=19). 5H. Relative proportion of TGFβR1 ligands in SAH liver samples. Outer ring shows the relative proportion of Scl1 (red colour) and other ligands of TGFβR1. 5I: Correlation plot showing correlation of Scl1 with clinical parameters.

We screened all proteins in secretome of *S. pyogenes* and *S. pneumonia* using mass spectrometry. Results showed certain collagen like proteins are present in the secretome. Based on literature evidence, it was known that *Streptococcus* species secretes surface proteins such as Streptococcal collagen-like protein 1 (Scl1), which exhibits structural homology to collagen; a canonical ligand of TGFβR1^25^.

Sequence alignment using BLAST between human collagen (NCBI accession: P02452) and Scl1 protein revealed a series of aligned, repetitive Gly-X-Y motifs (hallmarks of collagen-like sequences) indicating structural mimicry of Scl1 and collagen (Figure 5B, dot plot) suggesting a plausible mechanism by which Scl1 could act as a functional ligand for TGFβR1.

To predict the physical interaction between Scl1 and TGFβR1, molecular docking was performed using the HDOCK server. Among the top 10 predicted models, all showed a confidence score exceeding 80%, with model 1 showing the lowest docking energy and models 4 and 9 having minimal ligand RMSD values, signifying stable and high-affinity binding between Scl1 and TGFβR1 (Figure 5C and Figure 5D).

These results were experimentally validated using Co-immunoprecipitation mass spectrometry. Secretome from cultured *S. pyogenes* and *S. pneumonia* species were harvested and used to treat differentiated THP-1 macrophages. Co-immunoprecipitation-mass spectrometry analysis highlighted the presence of TGFβR1-Scl1 complexes, confirming a direct interaction (Figure 5D, Table S6). These results support the hypothesis that Scl1 secreted by *Streptococcus* species can bind to TGFβR1 and activate its downstream signalling.

We then evaluated the expression of Scl1 and *Streptococcus* in both stool and liver samples of SAH patients and ALD-rat models. Metaproteomic profiling revealed elevated levels of *S. pyogenes* and *S. pneumoniae* in stool samples of SAH patients (Figure 5E, Table S7). Scl1 levels were found to be elevated in ALD-rats and SAH→Healthy-rats and markedly reduced following FMT from healthy donor stool to ALD-rats (Figure 5F, Table S7). Scl1 abundance were also found to be significantly increased in stool samples of SAH patients (n=24, Table S4) compared to healthy controls (n=10) (Figure 5F, Table S7). This suggests that alcohol-associated dysbiosis fosters the expansion of *Streptococcus*, which in turn enhances Scl1 secretion.

Given that TGFβR1 is primarily expressed in the liver, we hypothesized that Scl1 translocates from the gut to the liver. Supporting this, Scl1 was detected at high levels in the liver tissues of ALD-rats and SAH stool treated rats, and SAH patients (n=19) compared to healthy liver. Importantly, human donor→ALD-rats showed reduced hepatic Scl1 levels, indicating that the presence of *Streptococcus* in the gut is directly linked to hepatic Scl1 abundance (Figure 5G, Table S7). Notably, Scl1 exhibited highest relative expression amongst all other potential ligands for TGFβR1 such as COL1A1, COL1A2, TGFβ1, and TGFβ2, in SAH liver highlighting that Scl1 could be a major contributor in activating TGFβ and associated downstream signalling in SAH patients (Figure5H).

In addition, Scl1 showed strong correlation with the severity indices in SAH patients (Figure 5I) suggesting its potential relevance in the pathophysiology of severe alcohol-related hepatitis and its possible association with disease progression.

### Therapeutic Impact of FMT on *Streptococcus* Abundance and Scl1 Levels in SAH patients

To further validate our results, stool and plasma samples from 16 SAH patients who underwent FMT were analysed at baseline and 28 days post-FMT (Figure 6A). Comparison of pre- and post-FMT stool samples revealed a significant reduction in 164 bacterial genera and an increase in 69 genera (p < 0.05) (Figure 6B). Notably, *Streptococcus* (genus level) was significantly reduced by 1.91-fold in stool samples collected 28 days after FMT (Figure 6C), which was accompanied by a reduction in plasma Scl1 protein levels at day 28 (Figure 6D, Table S8).

**Figure 6:**
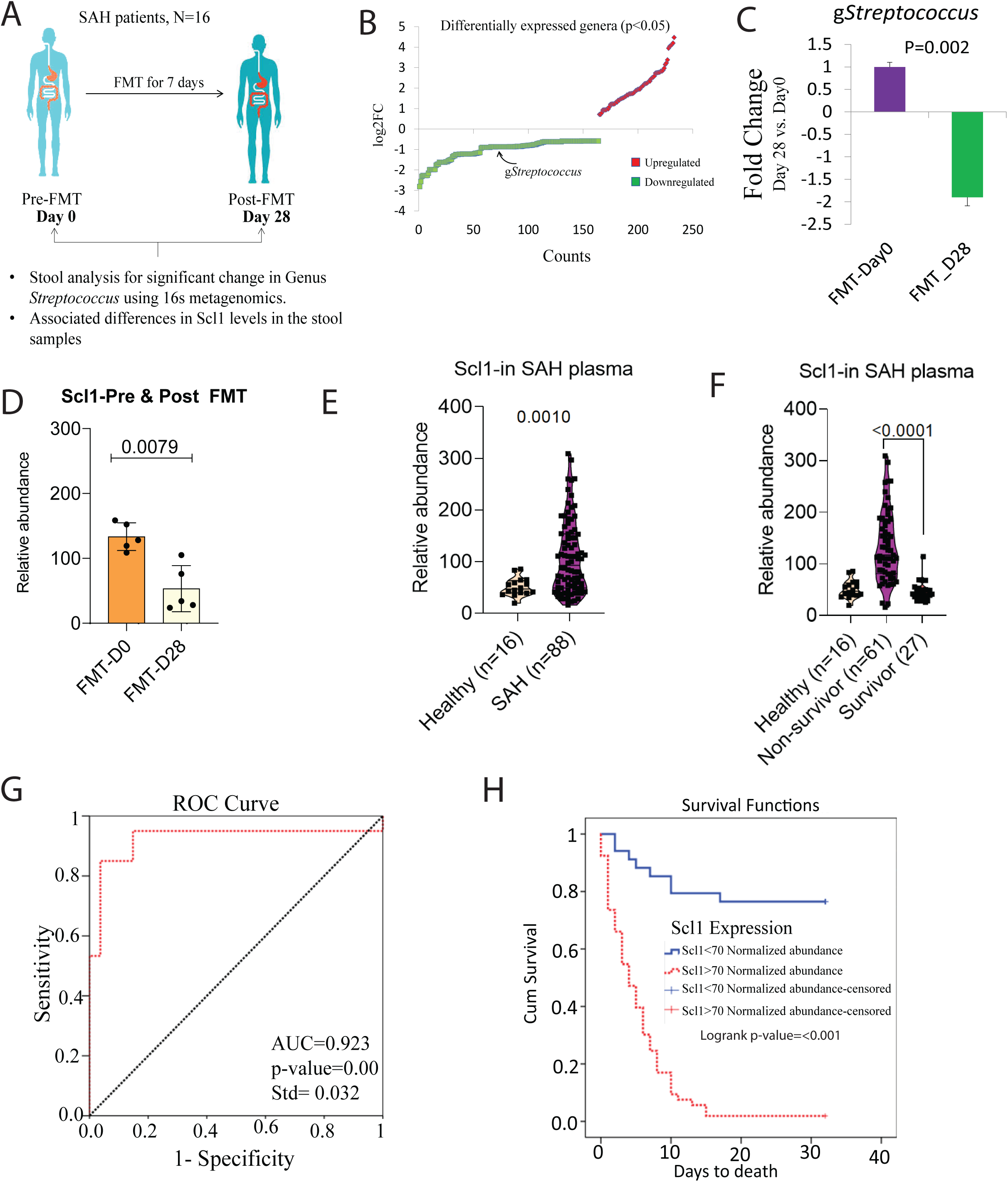
Therapeutic Impact of FMT on *Streptococcus* Abundance and Scl1 Levels in SAH patients: Figure 6A: Study design for patients who underwent FMT. Stool and plasma samples of 16 patients pre and post 28 days of FMT were analysed for *Streptococcus* and Scl1 levels Figure 6B: Differentially expressed genera in stool samples (post 28 day vs pre FMT). Figure 6C: Expression of g*Streptococcus* in stool of SAH patients (post 28-day vs pre FMT) Figure 6D: Expression of Scl1 protein in pre and post 28 day of FMT plasma samples. Figure 6E: Expression of Scl1 in SAH plasma (n=81) compared to healthy individuals (n=16). Figure 6F: Expression of Scl1 in SAH-survivor (n=27) and non-survivor (n=61) plasma and healthy individuals (n=16). Figure 6G: Receiver Operating Characteristic (ROC) Curve Analysis. The ROC curve (red dotted line) demonstrates the performance of the classifier in non-survivor. The curve shows high sensitivity and specificity with AUC of 0.923 and p-value 0.00. Figure 6H: Kaplan–Meier curves comparing overall survival between two patient groups. A cutoff level of >70 normalized abundance for Scl1 was selected to segregate survivor and non-survivor.

These findings collectively propose a novel gut-liver axis where alcohol-induced expansion of *Streptococcus* leads to elevated secretion of the collagen-mimic protein Scl1. Scl1 binds and activates hepatic TGFβR1 kinase (already known to be increased in ALD), driving pro-fibrotic and inflammatory signalling cascades that contribute to ALD pathogenesis.

To further establish Scl1 as a potential indicator of poor response and prognosis in severe alcohol-associated hepatitis (SAH), its plasma levels were quantified in healthy controls, SAH survivors, and non-survivors (demographics, Table S5). SAH patients exhibited significantly elevated plasma Scl1 levels compared to healthy controls (Figure 6E). Among SAH patients, non-survivors had markedly higher Scl1 expression than survivors, indicating its association with poor clinical outcome (Figure 6F). A normalized abundance cutoff of >70 effectively segregated non-survivors from survivors, with an AUROC of 0.92 and a p-value < 0.001, suggesting strong discriminatory power (Figure 6G).

Furthermore, Kaplan–Meier survival analysis demonstrated that patients with Scl1 levels above this threshold had significantly reduced 30-day survival compared to those with lower levels, confirming its potential as a predictive biomarker for short-term mortality in SAH (Figure 6H).

These findings highlight Scl1 as a promising prognostic biomarker in SAH, capable of predicting poor therapeutic response and 30-day mortality with high accuracy.

## Discussion

Gut dysbiosis has emerged as a critical determinant in the pathogenesis and progression of ALD^2,4,26^. Chronic alcohol consumption compromises the intestinal epithelial barrier, promotes microbial translocation, and disturbs the gut microbiota, favouring growth of pathogenic and pro-inflammatory bacterial species^27,28^. This dysbiosis shift facilitates systemic endotoxemia and propagates hepatic inflammation, immune activation, and fibrogenesis through the gut-liver axis^29^. While the downstream inflammatory and immunological effects of dysbiosis in ALD have been well studied, the precise microbial drivers and their mechanistic contributions modulating hepatic kinase signalling remain underexplored.

In this study, we provide compelling evidence that specific components of the alcohol-associated gut microbiota (AGM) can reprogram host liver kinase pathways to promote hepatic inflammation and fibrosis. We observed marked upregulation of kinases such as TGFβR1, BTK, CAMK4, MAP2K5, and IGF1R in the liver of ALD-rats and in rats transplanted with stool of SAH patients. These kinases are well-known regulators of inflammation, fibrosis, stress responses, and metabolic dysregulation-central features of progressive ALD^30^. Importantly, their expression was attenuated by FMT from healthy donors, underscoring the microbiota’s causative role in hepatic kinase reprogramming.

Stool meta-proteomic profiling revealed dysbiotic microbiome in both SAH patients and ALD-rats, dominated by abundance of *Streptococcus, Staphylococcus, Clostridium, Corynebacterium,* and *Mycoplasma*. These taxa have been previously associated with chronic liver injury and systemic inflammation ^31^. The reversal of activated kinases post-FMT with healthy human donor stool in rats showed a decline in the abundance of these bacteria, indicating that the pro-inflammatory and pro-fibrotic kinase signalling cascades are microbiota-dependent. Notably, 18 kinases, including BTK, MAP2K5, CAMK4, and TGFβR1, were significantly downregulated following FMT, suggesting a direct microbial influence on host kinase expression.

Of particular interest was the robust correlation between *Streptococcus* abundance and TGFβR1 expression (r=0.88), suggesting a potent and specific microbial trigger of the fibrogenic TGFβ axis. Additional associations with kinases such as BTK, MAP3K10, STK35, and CAMK1G strengthen the hypothesis that *Streptococcus* strains contribute to the inflammatory ^32^and fibrotic hepatic microenvironment in ALD/SAH. Supporting literature indicates that *Streptococcus* species are not only overrepresented in the ALD gut but these also secrete virulence factors capable of modulating host immunity and inflammation^32^.

In exploring the molecular mediator linking *Streptococcus* to TGFβR1 activation, secretome of *Streptococcus pyogenes* and *S. pneumoniae* were screened using proteomics. Among the identified proteins, several showed sequence similarity to collagen. Literature mining further revealed that Streptococcus secretes a collagen-like protein, Scl1, exhibiting the Gly-X-Y motif characteristic of mammalian collagen (Accession: P02452), a natural ligand for TGFβR1, thereby raising the possibility of functional mimicry. Previous studies have shown that Scl1 facilitates bacterial adhesion and immune evasion by mimicking host extracellular matrix components ^33,34^. Our findings expand its role to include receptor engagement and signal activation.

Using molecular docking, dot plot analysis, and sequence alignment, we confirmed the high-probability of interaction between Scl1 and TGFβR1. These computational predictions were validated experimentally, where the secretome of *S. pyogenes* and *S. pneumoniae* triggered TGFβR1 activation in THP1-derived macrophages. This interaction led to phosphorylation of SMAD2, activation of MAPK14 (p38), NFκB, IRF, and IFNαcanonical downstream effectors of TGFβ signalling, all of which were significantly elevated in liver of ALD-rats and SAH→Healthy-rats. These signals were substantially attenuated in HD→ALD-rats post-healthy human donor stool transplant, linking Scl1-driven receptor activation to the broader kinase reprogramming landscape observed in ALD.

Importantly, elevated Scl1 levels were confirmed in both the stool and liver tissues of SAH patients and ALD-rats, and these levels decreased following FMT. This supports the hypothesis that higher gut *Streptococcus* burden drives increased Scl1 production, which crosses the gut barrier to engage hepatic receptors. Such ligand mimicry by bacterial proteins introduces a paradigm shift in our understanding of host-microbe interactions in ALD from endotoxin-induced inflammation to direct microbial hijacking of host signalling pathways.

Beyond its ligand-mimetic role, Scl1 is implicated in several pathogenic processes. It promotes bacterial tissue invasion by binding integrins and extracellular matrix proteins, evades immune detection, and contributes to chronic infections and autoimmune-like inflammation^32–34^. These functions, coupled with its structural mimicry of collagen, suggest that Scl1 may provoke immune responses through molecular mimicry, a mechanism well-documented in autoimmune pathologies^35^. In ALD, where immune surveillance is already compromised, the translocation of Scl1 into systemic circulation represents a significant immune-modulatory threat.

Our findings also align with a growing body of literature demonstrating the role of bacterial proteins and metabolites in liver disease progression. For example, lipopolysaccharides (LPS) activate hepatic TLR4 signalling and drive inflammation and fibrogenesis^36^. Other microbial metabolites such as acetaldehyde, bile acid derivatives, and phenolic acids exacerbate oxidative stress and hepatocellular damage^37^. Proteins like pneumolysin (*S. pneumoniae*)^10^ and corisin^38^ (E. faecalis) have been shown to compromise epithelial integrity and induce profibrotic responses. Our identification of Scl1 adds to this expanding list of microbial effectors that disrupt host homeostasis and promote liver pathology.

To our knowledge, this is the first study to demonstrate that a bacterial product; Scl1 mimics human collagen and can directly activate fibrotic signalling pathways via TGFβR1 in the liver. This provides new insight into how gut microbes can subvert host cellular signalling in a manner analogous to cytokines or growth factors. It elevates the significance of microbial proteins as active effectors, capable of reprogramming host cell behaviour and contributing to chronic disease.

Clinically, our findings have several important implications. Targeting Scl1, whether by genetic ablation in bacterial strains, neutralizing antibodies, or small-molecule inhibitors, may offer a novel strategy for mitigating inflammation and fibrosis in ALD. Second, as Scl1 correlates directly with the severity scores, Scl1 could serve as a biomarker for disease progression or therapeutic response, particularly in patients with high *Streptococcus* abundance. Third, our results reinforce the therapeutic value of microbiota-targeted interventions such as FMT, not only for restoring microbial balance but for disrupting harmful microbial-host signalling interactions. Fourth, elevated plasma Scl1 levels were strongly associated with increased disease severity and poor clinical outcomes, including 28-day mortality, in SAH patients. Thus, Scl1 may serve as a predictive biomarker for short-term prognosis in severe cases.

However, several limitations warrant further investigation. We have not yet performed direct *in-vivo* inhibition of Scl1 or its interaction with TGFβR1. The mechanism of Scl1 translocation across the intestinal barrier remains to be elucidated. Moreover, while we demonstrated TGFβR1 activation in macrophages, future studies should evaluate its impact on hepatic stellate cells and hepatocytes to fully understand its fibrogenic potential.

To enhance translational relevance, future directions include inhibiting Scl1 in Streptococcus strains via genetic or antibody-based approaches in ALD models, and using decoy peptides or small molecules to block Scl1-induced TGFβR1 activation. Additionally, intravital imaging and biodistribution studies will trace Scl1 translocation from gut to liver, while investigations into antigen mimicry will assess its potential role in autoimmunity during chronic liver disease.

In summary, based on our results we identify Scl1;a collagen-mimetic bacterial protein secreted by *Streptococcus* species; as a novel microbial effector that activates hepatic TGFβR1 signalling and contributing to liver inflammation and fibrosis in ALD. These findings reshape our understanding of gut-liver interactions, highlighting the gut microbiota not merely as a source of inflammatory triggers but as an active modulator of host signalling.

The identification of Scl1 as a functional virulence factor that mimics host ligands to drive disease progression opens new avenues for diagnostics and microbiota-targeted therapies in ALD and potentially other fibrotic diseases.

## Supporting information

suplementary table all

## Abbreviations

ALD: Alcohol-related Liver Disease
SAH: Severe alcohol-related hepatitis
AST: Aspartate aminotransferase
ALT: Alanine aminotransferase
ROS: Reactive Oxygen Species
LDC: Lieber–De Carli
LE: Long Evan
mTOR: Mammalian Target of Rapamycin
MAPK14: Mitogen-Activated Protein Kinase 14 or p38
NFkB: Nuclear Factor-kappa B
IL6: Interleukin-6
TNFa: Tumor Necrosis Factor alpha
IL10: Interleukin-10

---

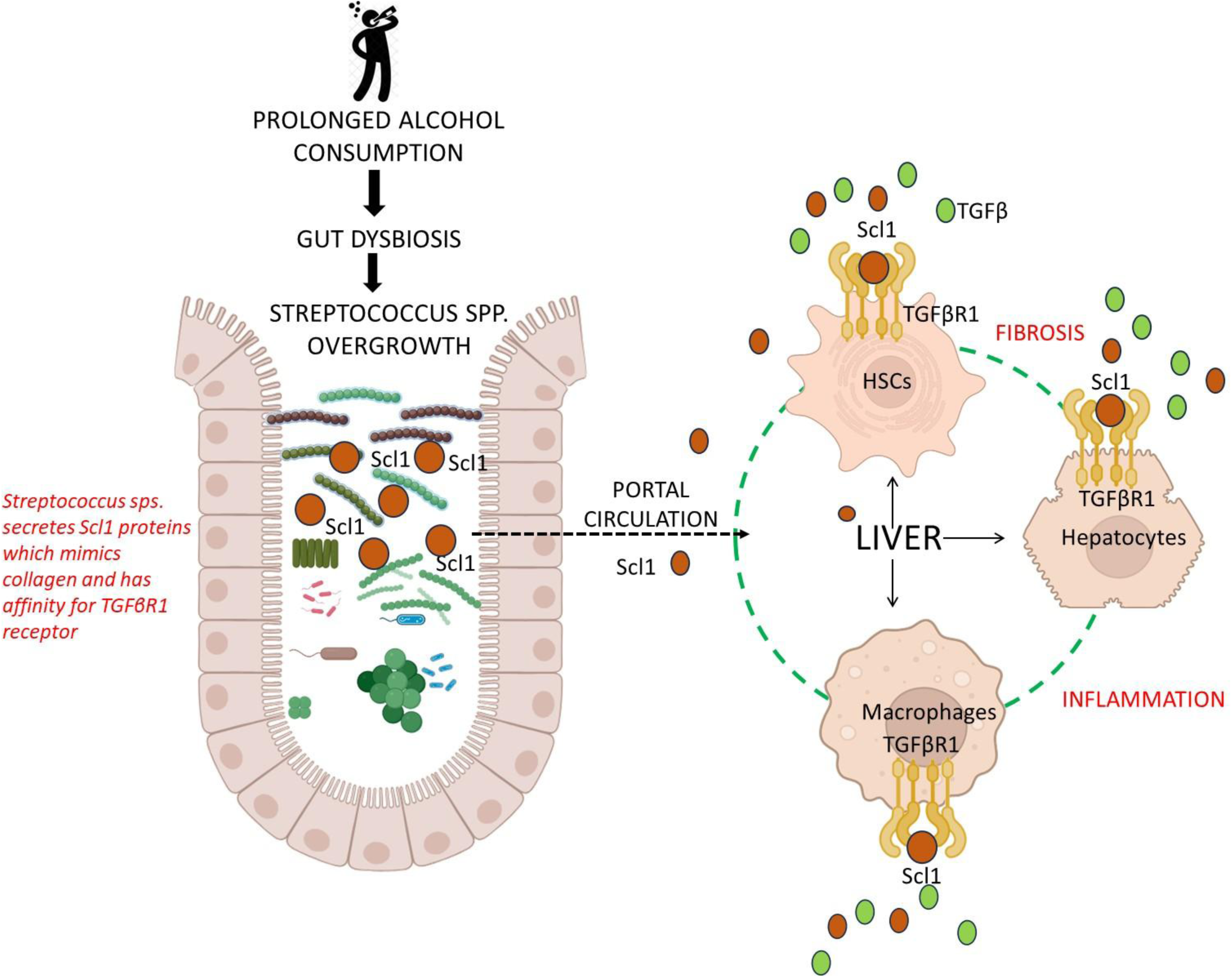

